# Cut and paste – Efficient cotyledon micrografting for *Arabidopsis thaliana* seedlings

**DOI:** 10.1101/751677

**Authors:** Kai Bartusch, Jana Trenner, Marcel Quint

## Abstract

Cotyledon micrografting represents a very useful tool for studying the central role of cotyledons during early plant development, especially their interplay with other plant organs with regard to long distance transport. While hypocotyl grafting methods are established, cotyledon grafting is still inefficient. By optimizing cotyledon micrografting, we aim for higher success rates and increased throughput in the model species *Arabidopsis thaliana*. We established a cut and paste cotyledon surgery procedure on a flat solid but moist surface which improved handling of small plant seedlings. Applying a specific cutting and joining pattern throughput was increased up to 40 seedlings per hour. The combination of short day conditions and low light intensities for germination and long day plus high light intensities and vertical plate positioning after grafting significantly increased ‘ligation’ efficiency. Together, we achieved up to 46 % grafting success in *A*. *thaliana*. Reconnection of vasculature was shown by successful transport of a vasculature-specific dye across the grafting site. On a whole plant level, plants with grafted cotyledons match plants with intact cotyledons in biomass production and rosette development. This cut and paste cotyledon-to-petiole grafting protocol simplifies the handling of plant seedlings in surgery, increases the number of grafted plants per hour and produces higher success rates for *A*. *thaliana* seedlings. The developed cotyledon micrografting method is also suitable for other plant species of comparable size.

## Introduction

Plant grafting has been successfully applied in various plant species for horticultural and agricultural purposes since ancient times [1–3]. Recently, grafting of small plant seedlings, often termed micrografting, has become an attractive tool to investigate physiological responses which depend on organ-to-organ long-distance transport of various substances [4]. Small transported molecules like plant hormones [5–8], small RNAs [9,10], small peptides/ proteins [11,12], nutrients [13,14], glucosinolates [15] and phytochelatins [16] have been already studied using micrografting.

Grafting in *Arabidopsis thaliana* was first described over 20 years ago [17] and was then utilized for different organs, including inflorescence [18], cotyledons [12] and shoots/roots [5,17]. While inflorescence micrografting and hypocotyl micrografting have reported success rates of up to 87 % [18] and 100 % [19,20], respectively, reported cotyledon micrografting (Cot-grafting) success rates have been very low (<2 %) so far [12]. Despite this constraint, Cot-grafting was already successfully applied to reveal the important role of cotyledons in floral development [12,21]. The mobile protein FLOWERING LOCUS T (FT) is produced in significant amounts in the cotyledons and transported via the phloem to the shoot apical meristem, where FT promotes floral induction [12].

Although the role of cotyledons in plant development and their interaction with other plant organs is of great importance [22–24], Cot-grafting has not been utilized for other approaches probably due to the low success rate of the described method [12]. Due to the size of the seedlings and their cotyledons, Cot-grafting is a technically challenging process, especially in *A*. *thaliana* and other small plant species. Frequently, the cotyledons do not remain attached to the petioles of the recipient plants, which is hypothesized to be caused by the oscillating movements of the petioles driven by the circadian clock [12]. This may be one of the main reasons for the low success rate.

As shoot-root hypocotyl grafting in *A. thaliana* has become a well-established method in fundamental plant research for various approaches [4], we aim to provide an optimized Cot-grafting method to encourage further research of cotyledons and their important role in plant growth and development. Therefore, we developed a flat-surface cotyledon-to-petiole grafting protocol which simplifies handling of grafted seedlings, increases the number of Cot-grafted plants per hour and produces high success rates (up to 46 %) for young *A. thaliana* seedlings.

## Methods

We developed Cot-grafting performed on a flat solid surface based on the general conditions for hypocotyl grafting [20]. The grafting procedure was then adjusted for cotyledon transplatation. Furthermore, the impact of different growth conditions before and after grafting on the success rate was evaluated.

### Plant material

Grafting experiments were performed with wild type *A. thaliana* seedlings from the Col-0 accession (obtained from the INRA collection as AV186). Using the growth conditions with the highest success rate (Table 1) for *A. thaliana*, Cot-grafting was also investigated in *Capsella rubella* (NASC ID N22697), *Arabidopsis suecica* (NASC ID N22505), *Brassica napus* (NASC ID N29003) and *Solanum lycopersicum* L. cv. Castlemart.

**Table 1.** The growth conditions before cotyledon micrografting (Cot-grafting) and the recovery conditions after Cot-grafting affect the graft union. Seedlings of *Arabidopsis thaliana* were grown at different light intensities, day length and grafted at different ages. Different light intensities, day length and plate positions were tested as recovery conditions after grafting. The experimenters KB and JT are indicated to evaluate human impact. 72 plant seedlings were grafted per combination and the graft union was evaluated 7 days after surgery. SD: Short day conditions (8 h of light and 16 h of darkness). LD: Long day conditions (16 h of light and 8 h of darkness).

| Growth conditions |  |  | Recovery conditions |  |  | Experimenter | Success rate (%) |
| --- | --- | --- | --- | --- | --- | --- | --- |
| Light ($\mu\text{mol}\cdot\text{s}^{-1}\cdot\text{m}^{-2}$) | Day length | Age (days) | Light ($\mu\text{mol}\cdot\text{s}^{-1}\cdot\text{m}^{-2}$) | Day length | Plate position | | |
| 30 | SD | 4 | 90 | SD | horizontal | KB | 36 |
|  |  |  |  |  | vertical | KB | 27 |
|  |  |  |  | LD | horizontal | KB | 42 |
|  |  |  |  |  |  | JT | 15 |
|  |  |  |  |  | vertical | KB | 46 |
|  |  |  |  |  |  | JT | 15 |
|  |  |  | 30 | LD | horizontal | KB | 14 |
| 90 | SD | 4 | 90 | LD | horizontal | KB | 17 |
|  |  |  | 30 | LD | horizontal | KB | 8 |
|  | LD | 4 | 90 | LD | horizontal | KB | 26 |
|  |  |  |  |  |  | JT | 6 |
|  |  | 5 | 90 | LD | horizontal | KB | 31 |
|  |  |  |  |  |  | JT | 21 |
|  |  | 6 | 90 | LD | horizontal | KB | 14 |
|  |  |  |  |  |  | JT | 22 |

### Growth conditions before Cot-grafting

In general, sterile conditions are recommended for micrografting [12,19,20]. Seeds were sterilized using a standard protocol [25] and stored at 4 °C for at least 7 days for stratification. Seeds were sown onto *A. thaliana* solution medium (ATS) [26] under a laminar flow hood. The plants were grown in a growth cabinet (Conviron Adaptis A1000) at 20 °C under different light and day length settings (Table 1): Long day (16 h light, 8 h darkness) and short day (8 h light, 16 h darkness) were applied as two different day lengths. The light intensity was either 30 or 90 µmol*s^−1^*m^−2^ (T5 white fluorescence lamps with 4000 K). The impact of the seedling’s age on Cot-grafting was investigated using 4-, 5- and 6-day-old plants.

### Cot-grafting procedure

The general micrografting environment described in detail by Melnyk [20] was implemented with some modifications: Although plant surgery is recommended to be carried out under a laminar flow, we worked on a lab bench under usual lab conditions. The dissecting tools – fine forceps and Ultra Fine Micro Knives (Fine Science Tools) – were sterilized with ethanol (70 %). All other materials were autoclaved before use, if possible. A Motic SMZ168 and a Zeiss Stemi DV 4 were used as dissecting microscopes. The general workflow of the developed Cot-grafting method and our established cutting patterns are illustrated in Figure 1 and 2. In brief, two round filter papers were moistened with distilled water and were placed into a sterile petri dish (Figure 1A). An excess of liquid was removed. Either one or two strips of Hybond N membrane were positioned on top of the filter papers (Figure 1B). Additional moistened strips of filter paper were used to maintain an optimal level of humidity during the grafting procedure. The recipient seedlings were transferred from the culture media plates to the nylon membrane using fine forceps (Figure 1C) and placed as flat as possible onto the membrane surface. A slight rotation of the hypocotyl could help to level out the seedling. Throughout the Cot-grafting procedure, it was very important to handle the seedlings carefully in order to prevent tissue damage. After these preparatory steps, the microsurgery was conducted under the dissecting microscope (Figure 1D). Both cotyledons of the recipient plant were removed by precise cuts using a microknife (recipient plant surgery – Figure 1F). Next, the cotyledons of the donor plant were cut off. A longitudinal cut along the central leaf vein was performed preparing the cotyledon and the petiole for optimal positioning to the recipient plant (donor cotyledon surgery; Figure 1G). The donor cotyledon was then transplanted to the recipient plant using forceps. If the cutting edges at the petioles did not fit well, a small piece of the petiole was cut off to obtain fitting angles between the petioles of the donor cotyledon and recipient plant. After surgery, the petri dishes were sealed using Parafilm® M.

**Figure 1.**
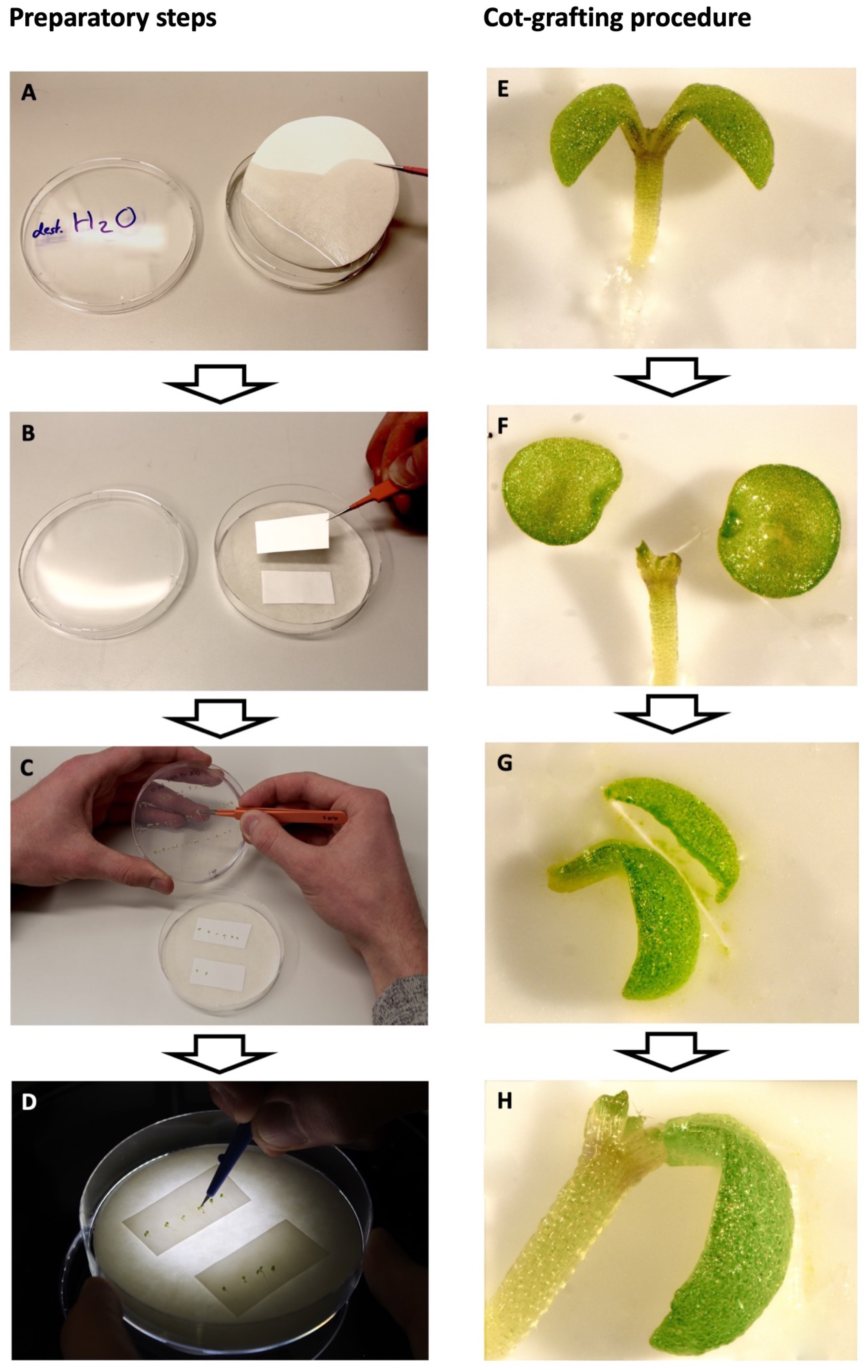
Workflow of cotyledon micrografting (Cot-grafting) preparation and procedure. A-D preparatory steps: Two layers of sterile filter paper were moistened with distilled water and placed into a sterile petri dish (A). Two stripes of nylon membrane were positioned on top of the filter papers (B). Vigorous seedlings were selected and placed onto the membrane using fine forceps (C). Cot-grafting was performed using a microknife and a dissecting microscope (D). E-H Cot-grafting procedure: The cotyledons of donor and recipient plants were cut off (E-F). The donor cotyledon was cut alongside the central leaf vein on one side for optimal positioning (G). The donor cotyledon was finally transferred to the recipient plant (H).

**Figure 2.**
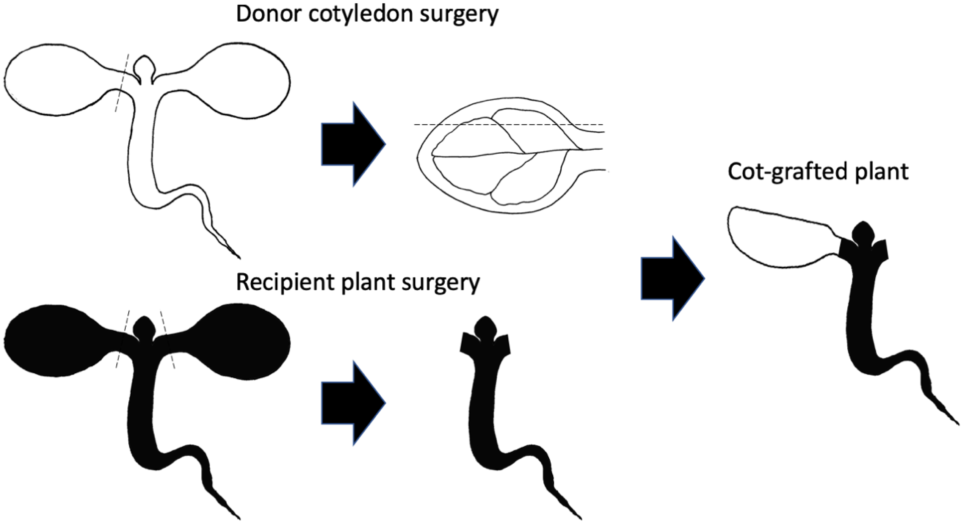
Cotyledon micrografting (Cot-grafting) surgery schematic. Donor cotyledon surgery: First, one cotyledon was cut from a donor plant. A second cut alongside the central leaf vein created a flat contact surface for optimal positioning. Recipient plant surgery: Both cotyledons of the recipient plant were cut off. The donor cotyledon was then transplanted to the recipient plant. Black dashed lines indicate cutting edges.

### Growth conditions after Cot-grafting

Cot-grafted plants were transferred back to the growth cabinet at 20 °C. Different recovery conditions after surgery were investigated (Table 1): The petri dishes were placed either vertically or horizontally. The light intensity was either 30 or 90 µmol*s^−1^*m^−2^ and day length was set to long or short day conditions.

### Evaluating the Cot-grafting success

To evaluate human impact on grafting success we also distinguished between the experimenters KB and JT. The transplanted cotyledons were evaluated 7 days after surgery using fine forceps. Well-connected cotyledons were counted as successful graftings. Detached or easily detachable cotyledons were counted as failed graftings. Apart from this, a vasculature specific dye, 5(6)-carboxyfluorescein diacetate (CFDA), was applied to identify reconnected vasculature as described by Melnyk et al (2015) [27] with the following modifications. For the evaluation of xylem connectivity, a piece of Parafilm® M was placed directly under the root of the Cot-grafted plant. 1 µl of 1 mM CFDA solution was dropped onto the root tip which was then cut for better dye penetration. The transported CFDA was detected in the cotyledons after 1 h using a stereomicroscope (Nikon SMZ1270) with GFP-filter and captured by a Nikon Digital Sight DS-Fi1c camera.

### Influence of Cot-grafting on plant development

The impact of the cotyledons on plant development was investigated by comparing biomass production of plants with one grafted cotyledon only and controls plants with two, one or without intact cotyledons. For germination and recovery conditions we used the combination with the highest grafting success rate: 4 days germination in 20°C, short day with a light intensity of 30 µmol*s^−1^*m^−2^ and 7 days recovery in vertical position in 20°C, long day with a light intensity of 90 µmol*s^−1^*m^−2^. The grafting procedure was carried out as described above and cotyledon removal of the control plants was done under the same conditions. After 7 days of recovery, all plants were transferred to ATS plates and kept under recovery conditions for additional 4 days. For the first biomass measurement, fresh weight of whole seedlings was recorded. Afterwards, the seedlings were transferred to soil and kept in the greenhouse for another 20 days. For the second biomass measurement, fresh weight of above ground material as well as the number of rosette leaves were recorded. The overall workflow is depicted in Figure 4A.

### Statistics

The collected data were statistically assessed by 1-way ANOVA followed by a Tukey HSD test (p < 0.05).

### Comparison of Cot-grafting methods

We applied the latest protocol about Cot-grafting [12] and compared the results to our method.

## Results and Discussion

### Success and limitations of Cot-grafting

By applying the established Cot-grafting protocol [12], we achieved a very low success rate of 1.25 %, similar to the published rates. In the following, we established the grafting conditions described in [20] and could also achieve success rates over 90 % in hypocotyl micrografting. We then developed a new cutting and joining approach for cotyledon grafting (Figure 1 and 2) and analyzed different combinations of conditions before and after grafting and their respective success rates (Table 1). The varying factor combinations resulted in grafting success rates ranging from 6 to 46 %. Most successful was the combination of short day and low light intensity before grafting and long day, high light intensity and vertical plate positioning during recovery. This combination lead up to 46 % grafting success and was then chosen for further experiments in this study.

Germination in short day conditions was reported to enhance the success rate in hypocotyl grafting [19]. We observed that despite lower grafting success (21 to 31 %) the handling of plants grown under high light and long day conditions before grafting was more efficient. The tissue of cotyledons and petioles was stronger and more vigorous under these conditions. Instead of 30 plants per hour using seedlings grown under low light and short day conditions, even up to 40 plants could be grafted per hour. Thus, the grafting success per time period is comparable to the plants of the factor combination with maximum success. We suggest that both growth conditions are justifiable.

Furthermore, as this is literally manual work, the impact of the experimenter should not be underestimated. While performing hypocotyl micrografting is known to be technically challenging [20], it was observed that Cot-grafting is probably even more difficult to perform due to the delicate petioles and cotyledons of young *A. thaliana* seedlings. In our case, success rates between experimenters differed. However, independent of the experimenter, success rates were consistently high.

Regarding the recovery conditions, high light intensity seems to have a positive influence on grafting success after surgery (Table 1). In contrast, the two tested day lengths for recovery as well as the mode of plate positioning did not affect grafting success rates.

An example of a successfully grafted *A. thaliana* cotyledon is depicted in Figure 3A. Xylem connectivity was successfully reestablished 7 days after grafting as shown by the detection of CFDA (Figure 3B). In addition, well-connected grafted cotyledons were sometimes still visible several weeks days after grafting (Figure 3C).

**Figure 3.**
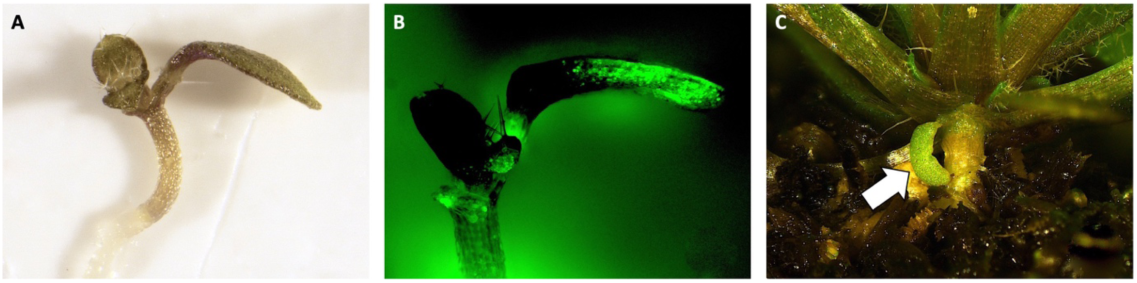
Evaluation of cotyledon micrografting (Cot-grafting) success. The Cot-graft junctions were evaluated 7 days after surgery. (A) Well-connected and vigorous cotyledons were counted as success. (B) Detection of 5(6)-carboxyfluorescein diacetate (CFDA) signal in the grafted cotyledon after application to the root tips indicates a successful xylem reconnection. (C) Well-transplanted cotyledons were still observable 31 days after grafting (arrow).

**Figure 4.**
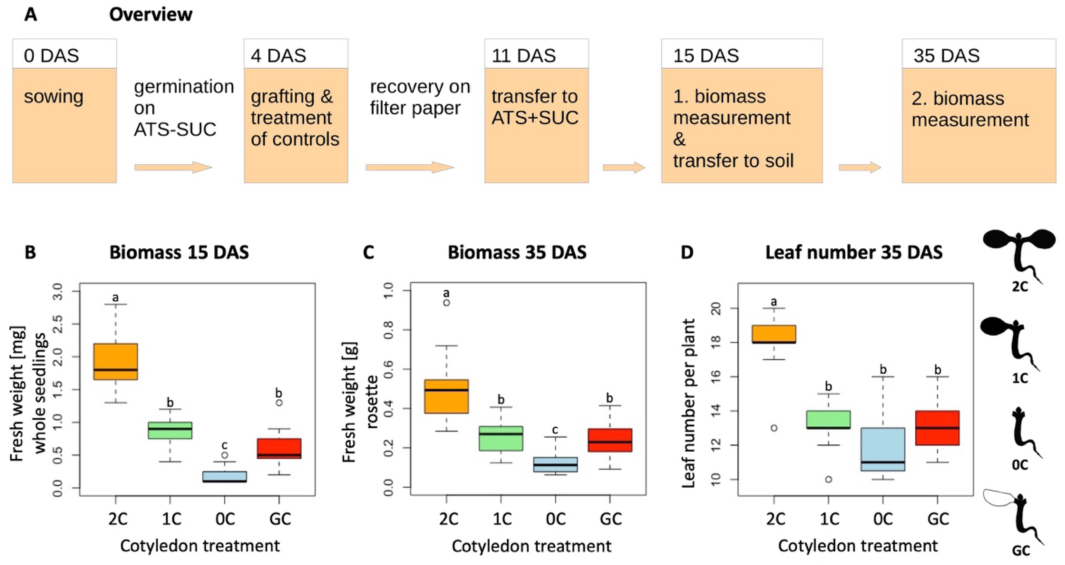
Grafted cotyledons and intact cotyledons affect biomass production equivalently. The impact of Cot-grafting on plant development was investigated by evaluating biomass production at two different developmental stages after grafting. (A) Illustration of the biomass evaluation workflow. (B-D) Fresh weight measurements of whole single seedlings 15 days after sowing (DAS; B). Above ground biomass of single plants 35 DAS (C), and number of rosette leaves 35 DAS (D). Cot-grafted plants (GC) were compared to three controls: intact plants with two cotyledons (2C), plants without cotyledons (0C) and plants with one cotyledon removed (1C). 15 plants were analyzed per cotyledon treatment. Different small letters denote statistically significant differences as assessed by 1-way ANOVA followed by a Tukey HSD test (p < 0.05). DAS: days after sowing, SUC: sucrose.

#### Successful replacement of cotyledons does not result in deficiency in plant growth

As cotyledons seem to have a huge impact on biomass production [23], a successful Cot-grafting method must not lead to deficiency in plant growth. Cot-grafted plants should have a comparable development to plants with intact cotyledons as shown in [23]. Therefore, we compared biomass production of plants with one grafted cotyledon only to the biomass of controls plants with two, one or without intact cotyledons.

At the time of the first biomass measurement, 11 days after grafting (15 days after sowing), control plants with two cotyledons had the highest single seedling fresh weight. Plants with one intact cotyledon showed a significantly lower fresh weight which was similar to the fresh weight of plants with one grafted cotyledon. The lowest fresh weight was measured for control plants without cotyledons (Figure 4B). This pattern was reproduced for the fresh weight measurements of the rosettes 31 days after surgery (Figure 4C). Meanwhile, only vital plants with two cotyledons showed a significantly higher leaf number than plants with only one or without cotyledons (Figure 4D). For all variants we observed that control plants with one intact cotyledon and plants with one grafted cotyledon perform similarly. Consequently, Cot-grafting seems to have no negative influence on plant growth.

#### Cot-grafting is applicable to other small plant species

For further practical tests, we applied our new Cot-grafting method to four other plant species. We chose *A. suecica* and *Capsella rubella* as similarly sized relatives of *A. thaliana. Brassica napus* and *Solanum lycopersicum* were chosen as crop plants of interest. We could not produce any successfully Cot-grafted plants in *Brassica napus* and *Solanum lycopersicum*. One or two days after grafting all transplanted cotyledons of these species were detached from the recipients and the hypocotyl had lost contact with the membrane. We hypothesize that the sturdier hypocotyls of *B. napus* and *S. lycopersicum* seedlings move more forcefully in their circadian oscillation [12] and detach easily from the moist surface. In contrast, well-connected cotyledons were observed in *A. suecica* (Figure 5A) and *C. rubella* (Figure 5B) with success rates of 9.5 and 1.9 %, respectively. There is, however, still room for improvement as we used the conditions optimized for *A. thaliana*. Thus, we find that this method is suitable for other small plant species. It could possibly be improved even for larger seedlings through a stabilizing tube (collar) across the graft junction as previously applied by Turnbull et al. [5] and Nisar et al. [18]. However, the conditions and probably even the procedure needs to be adjusted for each species separately.

**Figure 5.**
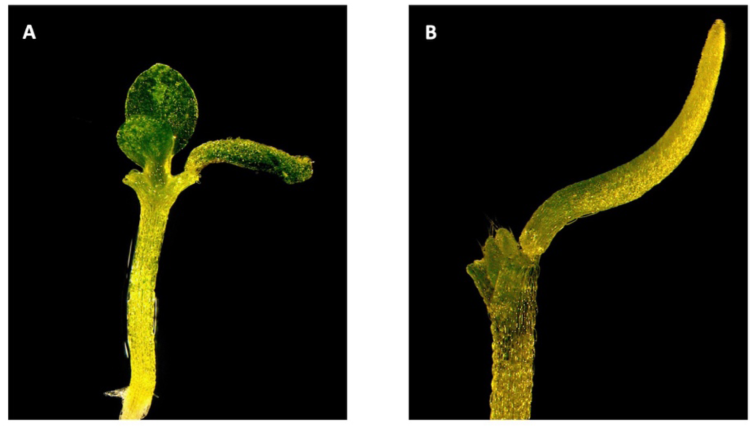
Cotyledon micrografting (Cot-grafting) is applicable to related plant species. Seedling of *Arabidopsis suecica* (A) and of *Capsella rubella* (B) with one grafted cotyledon. The seedlings recovered under long day conditions (16 h of light and 8 h of darkness) and 90 µmol*s^−1^*m^−2^ light intensity for 7 days.

Apart from this, cut-in grafting and hypocotyl-hypocotyl flat surface cutting between *A. thaliana* and *Eutrema salsugineum* was recently demonstrated by Li et al. [28]. Applying cut and paste cotyledon micrografting also to inter-species combinations would certainly expand the range of questions that could be addressed.

### Scion-to-petiole micrografting

Using the optimized Cot-grafting conditions, also scions can be transplanted from donor plants to petioles of recipient plants as illustrated in Figure 6A. Successfully scion-to-petiole grafted plants could be generated by this approach (Figure 6B and C), generating chimeric plants with two scions from different plants similar to the established Y-grafting method in the hypocotyl region [20]. This modification delivers another potentially interesting approach for joining plant organs coming from different origins to study organ-to-organ interactions.

**Figure 6.**
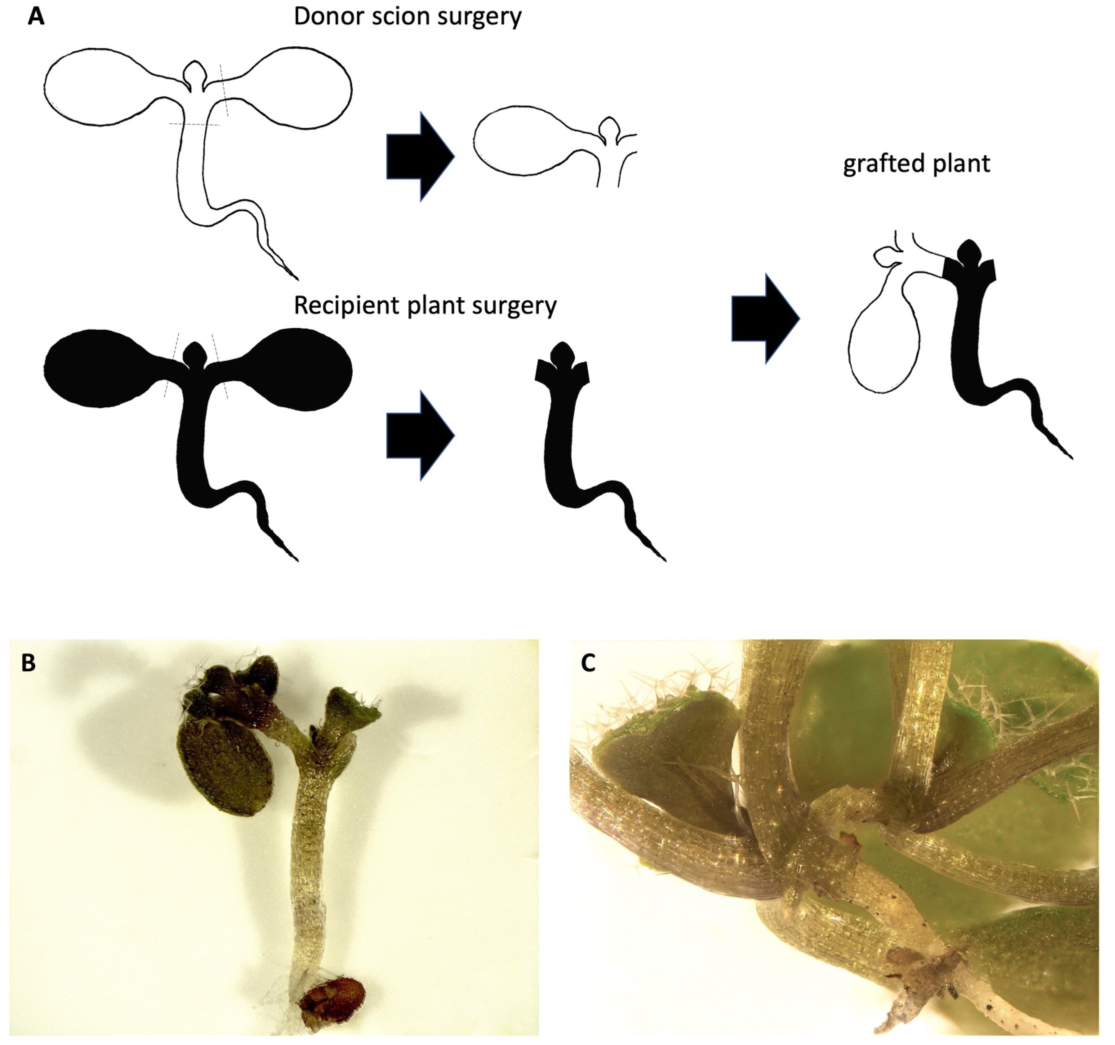
Scion-to-petiole micrografting. (A) The general procedure of Scion-to-petiole micrografting: the root and one cotyledon were removed from the donor plant first (donor scion surgery). Both cotyledons of the recipient plant were cut off (recipient plant surgery). The donor scion was then transferred to the recipient plant. Black dashed lines indicate cutting edges. A successful grafted plant 7 days (B) and 14 days after surgery (C).

## Conclusions

The success rate of Cot-grafting could be improved from 2 % up to 46 % through this modified method. A key step is to perform the grafting on a flat solid surface instead of grafting on cultural medium. Moreover, the adjusted cutting patterns could improve the efficiency of Cot-grafting. High light intensities after grafting are beneficial for the formation of the graft union. In general, Cot-grafting is a challenging process and requires practice, concentration and steady hands by the experimenter. The significant impact of cotyledons on biomass and leaf production during early plant development is maintained by successfully grafted cotyledons. In principle, Cot-grafting is possible in other plant species. However, it needs species-specific adjustment.

## Author contributions

KB and MQ conceived this project. KB, JT and MQ designed the study. KB and JT performed the experiments. KB and JT analyzed the data. KB and JT wrote the paper with contributions of MQ.

## Acknowledgements

We thank Kathrin Denk for her excellent technical assistance. We thank Rebecca Lippmann for capturing the pictures of Figure 1B-D and Syahfitri Wulandari for preparing the drawings of Figure 2, 4 and 6. We are grateful to Debora Gasperini and Adina Schulze who initially shared their expertise in hypocotyl grafting based on [20].

